# Progressive sub-MIC Exposure of *Klebsiella pneumoniae* 43816 to Cephalothin Induces the Evolution of beta-lactam Resistance without Acquisition of beta-lactamase Genes

**DOI:** 10.1101/2021.11.19.469034

**Authors:** Jasmine R. Anderson, Nghi B. Lam, Jazmyne L. Jackson, Sean M. Dorenkott, Taylor Ticer, Emir Maldosevic, Amanda Velez, Megan R. Camden, Terri N. Ellis

## Abstract

Bacterial exposure to antibiotic concentrations below the minimum inhibitory concentration (MIC) may result in a selection window allowing for the rapid evolution of resistance. These sub-MIC concentrations are commonly found in the greater environment. This study aimed to evaluate the adaptive genetic changes in *Klebsiella pneumoniae* 43816 after prolonged but increasing sub-MIC levels of the common antibiotic cephalothin over a fourteen-day period. Over the course of the experiment, antibiotic concentrations increased from 0.5 μg/mL to 7.5 μg/mL. At the end of this extended exposure, the final adapted bacterial culture exhibited clinical resistance to both cephalothin and tetracycline, altered cellular and colony morphology, and a highly mucoid phenotype. Cephalothin resistance exceeded 125 μg/mL without the acquisition of beta-lactamase genes. Whole genome sequencing identified a series of genetic changes that could be mapped over the fourteen-day exposure period to the onset of antibiotic resistance. Specifically, mutations in the *rpoB* subunit of RNA Polymerase, the *tetR/acrR* regulator, and the *wcaJ* sugar transferase each fix at specific timepoints in the exposure regimen where the MIC susceptibility dramatically increases. These mutations indicate that alterations in the secretion of colanic acid and attachment of colonic acid to LPS, may contribute to the resistant phenotype. These data demonstrate that very low, sub-MIC concentrations of antibiotics can have dramatic impacts on the bacterial evolution of resistance. Additionally, this study demonstrates that beta-lactam resistance can be achieved through sequential accumulation of specific mutations without the acquisition of a beta-lactamase gene.

**Importance:** Bacteria are constantly exposed to low levels of antibiotics in the environment. The impact of this low-level exposure on bacterial evolution is not well understood. In this work, we developed a model to expose *Klebsiella pneumoniae* to progressive, low doses of the antibiotic cephalothin. After a fourteen-day exposure regimen, our culture exhibited full clinical resistance to this antibiotic without the traditional acquisition of inactivating genes. This culture also exhibited resistance to tetracycline, had a highly mucoid appearance, and exhibited altered, elongated cellular morphology. Whole genome sequencing identified a collection of mutations to the bacterial genome that could be mapped to the emergence of the resistant phenotype. This study demonstrates that antibiotic resistance can be achieved in response to low level antibiotic exposure and without the traditional acquisition of resistance genes. Further, this study identifies new genes that may play a role in the evolution of antibiotic resistant bacteria.

## Introduction

The mis- and overuse of antibiotics by the medical and agricultural industries has created a basal level of antibiotic exposure present in our environment (1–5). Antibiotics may be introduced into the environment via three major pathways. The first is from urine and excretions of people, pets, and livestock. Between 40-90% of administered antibiotics are excreted through urine and feces with the molecule still in its active form (1). Second, antibiotics such as chlorhexidine are used in commercial agriculture and aquaculture (3). These substances can then contaminate nearby lands via wastewater and groundwater seepage (1, 6). Finally, antibiotics can enter the local environment through improper disposal of unused or expired prescriptions (1, 3). While concentrations of antimicrobial substances in the environment will vary based on location and potential sources of contamination, it is generally agreed that the residual environmental antibiotic concentration is not high enough to eradicate native bacterial populations. However, even at low concentrations these compounds become a source of survival stress for bacteria in the soil and water supplies. Concentrations below the bacteriostatic limit, referred to as sub-MIC levels, may result in the creation of a selection window for bacteria (7–9). This window is the concentration at which the occurrence of genomic mutations is highest and can lead to the development of clinical antibiotic resistance in pathogens.

Experiments have shown that sub-MIC antibiotic treatment can alter the resistance profile, nutrient use, protein expression, gene transcription and mutation rates among common nosocomial pathogens (4, 10, 11). However, current studies of sub-MIC exposure and bacterial adaptation have a number of limitations. First, these experiments frequently utilize strains of bacteria either known to harbor antibiotic resistance genes or are otherwise clinically resistant to one specific class of antibiotic. Second, the experimental designs include only brief exposure times to antibiotics. In most studies, the samples were exposed to the antibiotic for only 24-48 hours, providing limited time for the bacteria to evolve novel genetic changes like those seen in a clinical or environmental setting where antibiotic exposure is both consistent and long term. Finally, genetic analyses frequently used pre-determined targets, or the phenotypes assumed from mutations in predetermined targets. While the results of these studies are informative, they are not comparable to situations in which antibiotic resistance phenotypes evolve over time through prolonged sub-lethal antibiotic exposure.

In this study, we utilized *Klebsiella pneumoniae* 43816, which does not exhibit clinical resistance to the major antibiotic classes. Bacterial cells were cultured using a progressive exposure method to gradually increasing sub-MIC concentrations of the beta-lactam antibiotic cephalothin over a fourteen day period. At the end of this prolonged exposure period to sub-MIC concentrations the bacteria exhibited clinical resistance to both cephalothin and tetracycline, altered cellular and colony morphology, and a highly mucoid phenotype. Whole genome sequencing identified a series of genetic changes that could be mapped over the fourteen-day exposure period to the onset of antibiotic resistance. Specifically, mutations in the *rpoB* subunit of RNA Polymerase, the *tetR/acrR* regulator, and the *wcaJ* sugar transferase each fix at specific points in the exposure regimen where the MIC susceptibility dramatically increases. These mutations indicate that secretion and export of colanic acid, and its association to lipopolysaccharides, may contribute to the resistant phenotype. These data demonstrate that very low, sub-MIC concentrations of antibiotics can have dramatic impacts on the bacterial evolution of resistance. Additionally, this study demonstrates that antibiotic resistance can be achieved through accumulation of mutations without the traditional acquisitions of beta-lactamase or other drug inactivating genes.

## Materials and Methods

### Bacterial Strains and Progressive Antibiotic Exposure

*Klebsiella pneumoniae* 43816 (ATCC, Manassas, VA) was used as the starting culture for the progressive antibiotic exposure experiment (Figure 1). All cultures were grown in Luria-Bertani (LB) broth (BD Difco, Franklin Lakes, NJ) at 37°C with shaking at 200 rpm. All antibiotics and reagents are from Thermo Fisher Scientific (Waltham, MA) unless otherwise indicated. *K. pneumoniae* was grown overnight from frozen stocks and 50 μL of this culture was added to 5 mL of fresh LB with 0.5 μg/mL cephalothin added. The concentration of 0.5 μg/mL cephalothin was significantly below the MIC for this organism. Cultures were grown for 12 hours, at which point 50 μL were transferred to 5 ml of fresh LB with the same concentration of cephalothin. After a total of 24 hours exposure to one dose of antibiotics, 50 μL of culture were transferred to a new 5 mL of LB with a higher concentration of cephalothin. Each stepwise exposure increased the dose of cephalothin by 0.5 μg/mL, to a final dose of 7.5 μg/mL cephalothin. An untreated culture of *K*. *pneumoniae* 43816 was grown in parallel without the addition of antibiotic. Frozen glycerol stocks were made of each culture at each 12-hour transfer point for later analysis. *Klebsiella pneumoniae* 43816 Δ*wcaJ* (VK646) was kind gift from Kimberly A. Walker and Virginia Miller at UNC Chapel Hill (12).

**Figure 1:**
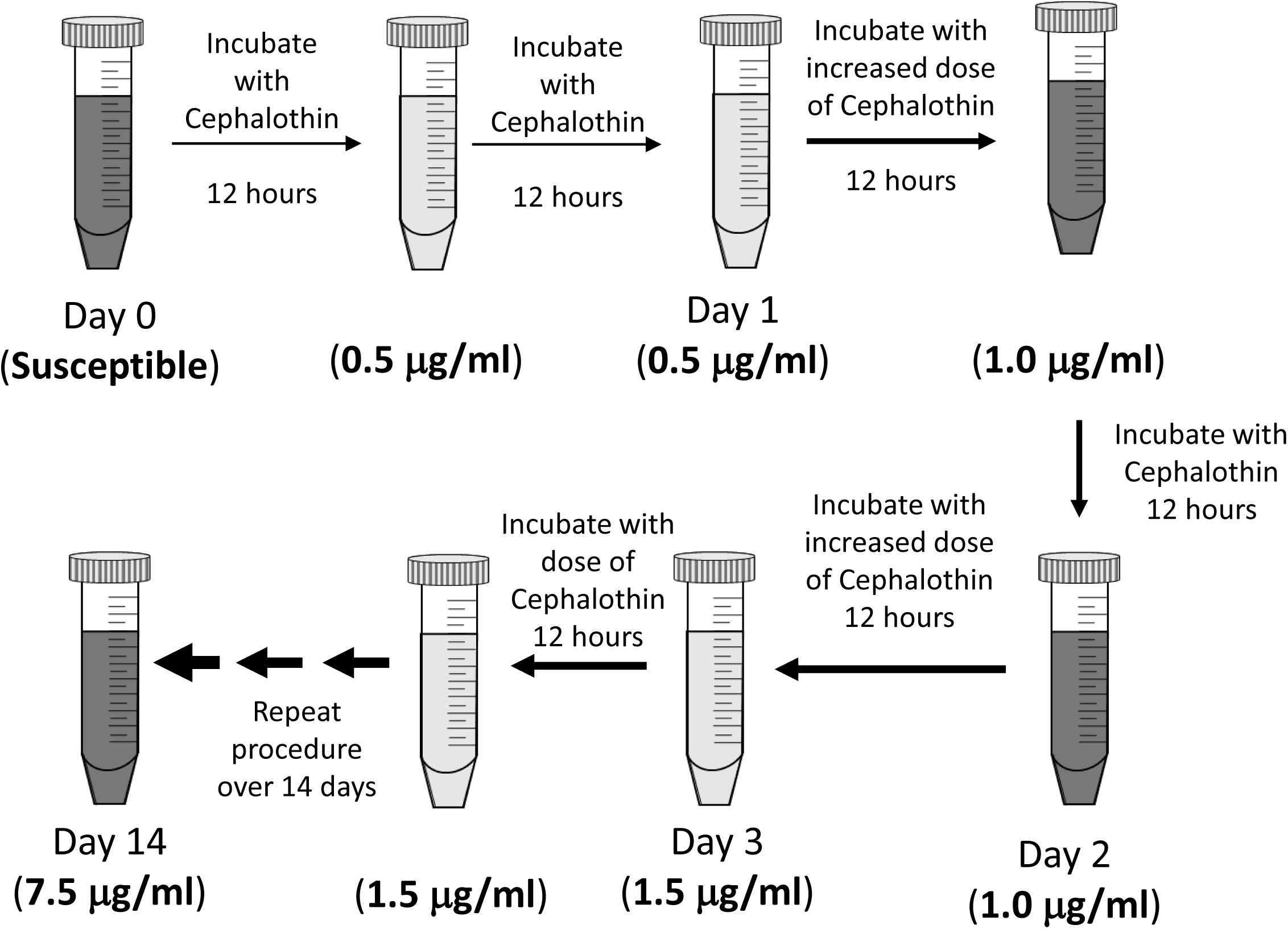
Experimental design. *Klebsiella pneumoniae* 43816 was exposed to increasing concentrations of cephalothin in LB media over a continuous 14-day period. All cultures were grown in LB media at 37C with 200 rpm shaking. Each single dose exposure lasted 24 hours and included a sub-culture into fresh media with antibiotics at the 12-hour mark. Frozen bacterial stocks were made every 12 hours for subsequent analysis. A control culture was also grown, sub-cultured, and sampled in LB media without antibiotics following the same schedule as the treated sample.

### Bacterial Growth and Morphology

At the 12-hour timepoint in each exposure, the bacteria were transferred to fresh LB with antibiotic. At this point, the time required for the bacterial culture to reach an OD600 of 1.0 was determined using an Eppendorf Biophotometer (Hamburg, Germany). Decreases in the growth rate were determined by subtracting the elapsed time for the treated culture from the average time to OD600 1.0 of all untreated cultures (142 minutes).

Changes in colony morphology were determined by quadrant streaking on LB agar plates. To determine cellular morphology, overnight bacterial cultures were centrifuged at 5,000 rpm for 5 minutes and re-suspended in phosphate-buffered saline (PBS) (pH 8.0). Samples were then diluted in PBS to an optical density of approximately 1.0 OD600. Cells were negative stained with 1% Nigrosin and visualized by brightfield microscopy at 1,000 x with an oil immersion lens.

### Determination of Minimum Inhibitory Concentration Breakpoints

Broth microdilution assays were used to determine the minimum inhibitory concentration (MIC) of tetracycline, amikacin and five different beta-lactam antibiotics (Cephalothin, Cefoxitin, Cefotaxime, Cefepime, and Imipenem). The beta-lactam drugs were chosen as representative of different generations of cephalosporin antibiotics and a carbapenem, respectively. 96 well plates were seeded with LB broth containing 10^3^ CFU/mL starting concentration of bacteria. Antibiotics were added in a two-fold serial dilution starting at 500 μg/mL. Cultures were incubated overnight, and growth was measured at OD600 using a Biotek Gen5 plate reader (Winooski, VT). Wells containing only LB and bacteria were used as controls for normal growth. The MIC breakpoint was determined as the lowest antibiotic concentration at no increase in optical density over a broth only control was detected. All samples were tested in triplicate.

### Genomic DNA Isolation

Genomic DNA was extracted using a protocol modified from Wright et. al. (13). Briefly, 50 mL of overnight culture was pelleted for 10 minutes at 10,000 rpm at 4°C. The pellet was washed twice with TE25S buffer (25 mM Tris-HCl, 25 mM EDTA, 0.3 M sucrose, pH 8.0) and resuspended in TE25S with 10 mg/mL lysozyme and RNAse. The mixture was incubated for two hours at 37°C with shaking at 150 rpm. Proteinase K and 10% SDS were added and incorporated by inversion and incubated for 1-2 hours at 50-55°C with periodic inversions. 5M NaCl was added followed by 3.25 mL of CTAB (Cetyl Trimethyl Ammonium Bromide)/NaCl. The solution was mixed by inversion and incubated at 55°C for 10 minutes. A 24:1 chloroform/isoamyl alcohol solution was added and incubated at room temperature with shaking at 100 rpm for 20 minutes. After incubation, the solution was centrifuged at 10,000 rpm and 4°C for 15 minutes, and the upper aqueous layers were transferred into fresh tubes. This chloroform treatment was repeated a second time. The upper aqueous layers of both samples were combined with an 0.6 volume of isopropyl alcohol and the mixture was gently inverted. After five minutes, the purified DNA was spooled from the tube onto a sterile Pasteur pipette. The spooled DNA was washed with approximately 5 mL of 70% ethanol and dried before being suspended in 300µL of EB buffer (QIAGEN, Hilden, Germany). Purified DNA was quantified by Qubit (Thermofisher).

### Genome Sequencing

Whole genome sequencing was performed on the following *K. pneumoniae* samples: days 1-14 of the adaptation experiment the small colony forming variant, the large colony forming variant, and the untreated sample after 14 days of culture. These DNA samples were analyzed by the Microbial Genome Sequencing Center (https://www.migscenter.com) which utilized an Illumina sequencing technique similar to that used by Baym *et. al*. (14). Samples were compared against the published reference sequence NZ_CP009208.1 for *K. pneumoniae* 43816 (15). Any variations were analyzed using a proprietary breseq variant calling algorithm (16).

To confirm these results, and to extend our analysis to include SNPs within promoter regions, endpoint samples of DNA from *K. pneumoniae* 43816, the fourteen-day antibiotic adapted culture, and the fourteen-day untreated population were also analyzed using SMRT Cell Sequencing by the National Center for Genome Resources (www.ncgr.org). Samples were again compared against the published reference sequence NZ_CP009208.1 for *K. pneumoniae* 43816 (15). Genetic variation between genomes was calculated using a modified form of FST or analysis of variance referred to as ϴ (17). FST values are evaluated against the null hypothesis that the populations are not genetically unique (17). Pacific Biosciences calculated allele frequencies and utilized a proprietary Quiver Algorithm to maximize accuracy in sequence reads using variation between the published genome and prior records.

The complete genome sequences are available via the NCBI BioProject data base. The project ID is PRJNA854906.

### Capsule Extraction and Characterization

Capsular polysaccharide (CPS) was extracted using the protocol outlined by Domineco *et al*. (18). Briefly, 500 μL of an overnight culture were mixed with 100 µL of 1% Zwittergent 3-14 in 100 mM citric acid, pH 2.0. The mixture was vortexed vigorously, incubated at 50°C for 20 minutes, and centrifuged for 5 minutes at 14,000 rpm. The supernatant was then transferred to a fresh tube, mixed with 1.2 mL of absolute ethanol, and incubated for 90 minutes at 4°C. Precipitate was collected after centrifugation at 14,000 rpm for 10 minutes and dissolved in 200 µL DI water.

CPS D-glucuronic acid was quantified following a previously established protocol by Lin *et al.*(19). Purified CPS was vortexed vigorously with 1.2mL of 12.5mM sodium tetraborate in concentrated sulfuric acid and heated for 5 minutes at 95°C. The samples were cooled before the addition of 20 µL of 0.15% m-hydroxydiphenyl; and the absorbance was measured at 540 nm. A standard curve was generated using D-glucuronic acid to determine the concentration of glucuronic acid in the CPS samples. To ensure quantification of CPS from the same number of bacteria, strains were normalized to 10^8^ CFUs/mL. Each assay was performed in triplicate from six individual cultures.

The extracted capsule samples were also quantified for protein content and sialic acid content. Protein content was determined using a BCA Protein Assay kit (Thermo Scientific), according to manufacturer’s directions. Sialic Acid was quantified using the Sialic Acid Quantitation Kit (Sigma), according to manufacturer’s directions.

The carbohydrate composition of the CPS was characterized by GC-MS as previously detailed in Brunson et al (20). Briefly, CPS was purified from LPS using sodium deoxycholate at a final concentration of 6 mM as described by Kachlany et al. (21). The CPS was pelleted with cold ethanol, then freeze-dried before being hydrolyzed in 0.5 M HCl at 85°C for 18 h. The hydrolyzed carbohydrates were modified with the Tri-Sil HTP reagent (Thermo Scientific, Wltham MA.), as described by York et al. (22). The modified carbohydrates were dried and resuspended in 1 mL hexane. The carbohydrate suspension in hexane was centrifuged at 1000 x g for 5 minutes and the supernatant was collected for GC-MS analysis.

GC-MS analyses were conducted with a CP-3800 GC (Varian, Palo Alto, CA.) using a Supelco SPB-608 30-m fused silica capillary column, containing a bonded stationary phase (0.25 μm film thickness). The TMS (tri-methyl silyl) conjugated glycans were analyzed using the electron ionization mode with a Saturn 2200 GC/MS (Varian, Palo Alto, CA.). The initial oven temperature was 80 °C, held for 2 min. Then, the temperature was raised by 20°C/min and held at 160°C for 12 min. Finally, the oven temperature was raised by 20°C/min and held at 260°C for 7 min.

### Quantification of Lipopolysaccharide

Whole cell lipopolysaccharide levels were quantified using the Purpald assay (23, 24). Briefly stated, cultures were centrifuged at 5,000 rpm for 5 minutes and re-suspended in PBS. 50 µL of this cell solution was treated with 32 mM sodium periodate solution in a 96-well plate and incubated at room temperature for 25 minutes. After this incubation period, a 136 mM Purpald solution in 2N NaOH was added, and the plate was incubated for an additional 20 minutes. Following the completion of this step, 64 mM sodium periodate solution was added, and the plate was incubated for 20 minutes. Absorbance was immediately determined at 540 nm and compared to a standard curve of pure lipopolysaccharide isolated from *K. pneumoniae* (Sigma. St. Louis MO.). This assay was also used for capsule extracted as described above for the glucuronic acid assay. Results are presented with LPS concentrations normalized to 10^8^ CFUs.

### Statistical Analysis

Statistical significance was determined using a one-way ANOVA and Tukey’s post hoc test using XLSTAT software. Significance for the D-glucuronic acid and protein quantification was compared against results from the time zero, untreated sample. Significance was determined at a p value of ≤0.01 or less.

## Results

### Experimental design of progressive sub-MIC exposure to cephalothin

This study aimed to evaluate the adaptive genetic changes in *Klebsiella pneumoniae* 43816 upon prolonged but increasing exposure to sub-MIC levels of the common antibiotic cephalothin. *Klebsiella pneumoniae* 43816 was utilized because *Klebsiella* is known for its genomic plasticity and rapid evolution of antibiotic resistance (25, 26). Additionally, the genome of this strain has been sequenced (15), which allows for genetic changes to be tracked over the course of the progressive exposure.

The experimental design of this exposure protocol is diagramed in Figure 1. Fresh cultures were grown in LB at 37C with 200 rpm shaking and an initial cephalothin concentration of 0.5 μg/mL. After the first 12 hours of culture, a sample was transferred to fresh LB with the same antibiotic concentration and allowed to grow for a second 12-hour period. The bacteria were exposed to any one dose of antibiotics for 24 hours in total. Then, the sample was transferred to a culture with an increased concentration of antibiotic. At each step, the antibiotic concentration was increased by 0.5 μg/mL, to a final exposure of 7.5 μg/mL, which was well below the CLSI standard of 16 μg/mL for cephalothin resistance (27). As a control, an untreated culture, without antibiotics, was grown and transferred in parallel to LB without any antibiotic. Frozen stocks were made of all cultures at each 12-hour timepoint and saved for later analysis.

### Progressive antibiotic exposure alters *K. pneumoniae* growth, cellular morphology, and colony morphology

In order to evaluate the impact of such low antibiotic concentrations, we measured the rate of growth of the culture at the 12-hour transfer mark. This was determined by measuring the time it took from the culture to reach an optical density (OD600) of 1.0 after subculturing. As seen in Table 1, the cultures exposed to antibiotics exhibited slower growth (time to OD 1.0), which is indicative of cellular stress. The growth rate of the treated culture was substantially slowed once antibiotic concentrations exceeded 5 μg/mL; while the growth rate of the untreated culture was not significantly affected over the entire course of the experiment.

**Table 1:**
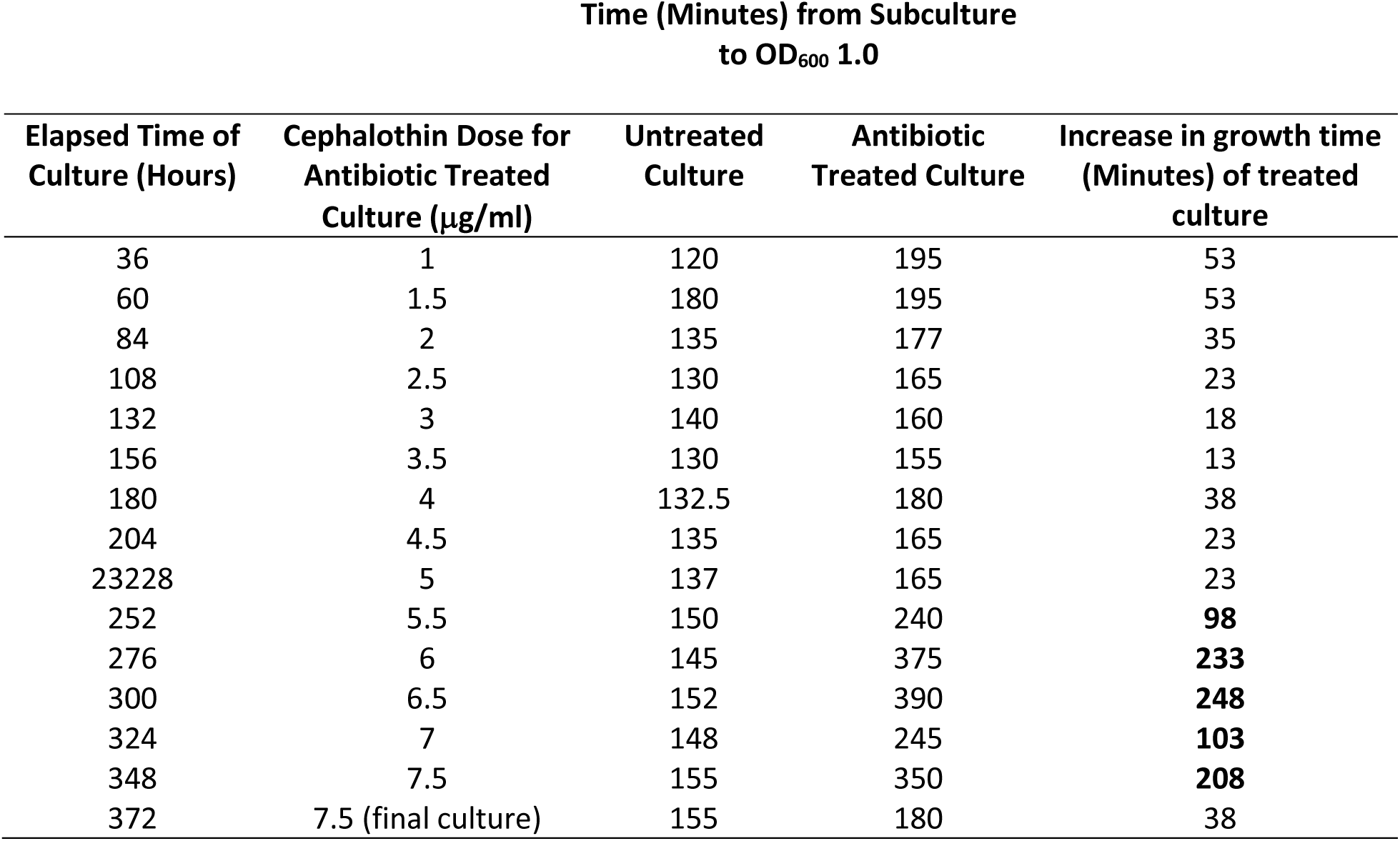
Changes in growth rates because of low dose antibiotic exposure. The initial impact of sub-MIC antibiotic exposure was determined by evaluating the growth rate of cultures upon transfer to a new antibiotic concentration. Measurement of time required for adapted and untreated *K. pneumoniae* 43816 sub-culture to reach an OD600 of 1.0 was measured after 12-hour exposure at each time point/dose. Changes in growth between treated and control was determined relative to the average growth time for all untreated cultures (142 minutes). Values in bold represent increases in growth time greater than one half that of the untreated culture.

This slow growth rate was not a permanent effect, as demonstrated by the growth properties of the final adapted culture (Table 1). The final adapted culture was able to grow rapidly in 7.5 μg/mL cephalothin (time to OD600 1.0). This was similar to the times recorded for the untreated culture at any time point during the fourteen days of continuous growth in LB.

While the growth rates of the adapted culture returned to a range similar to the untreated culture, the appearance of the antibiotic treated culture was significantly and permanently altered after the progressive exposure experiment. The antibiotic adapted culture exhibited a highly mucoid appearance in liquid culture when compared to the clonally related untreated group (Figure 2A). While the viscosity changed, the adapted population did not qualify as a hypermucoviscous strain as indicated by a string test (data not shown).

**Figure 2:**
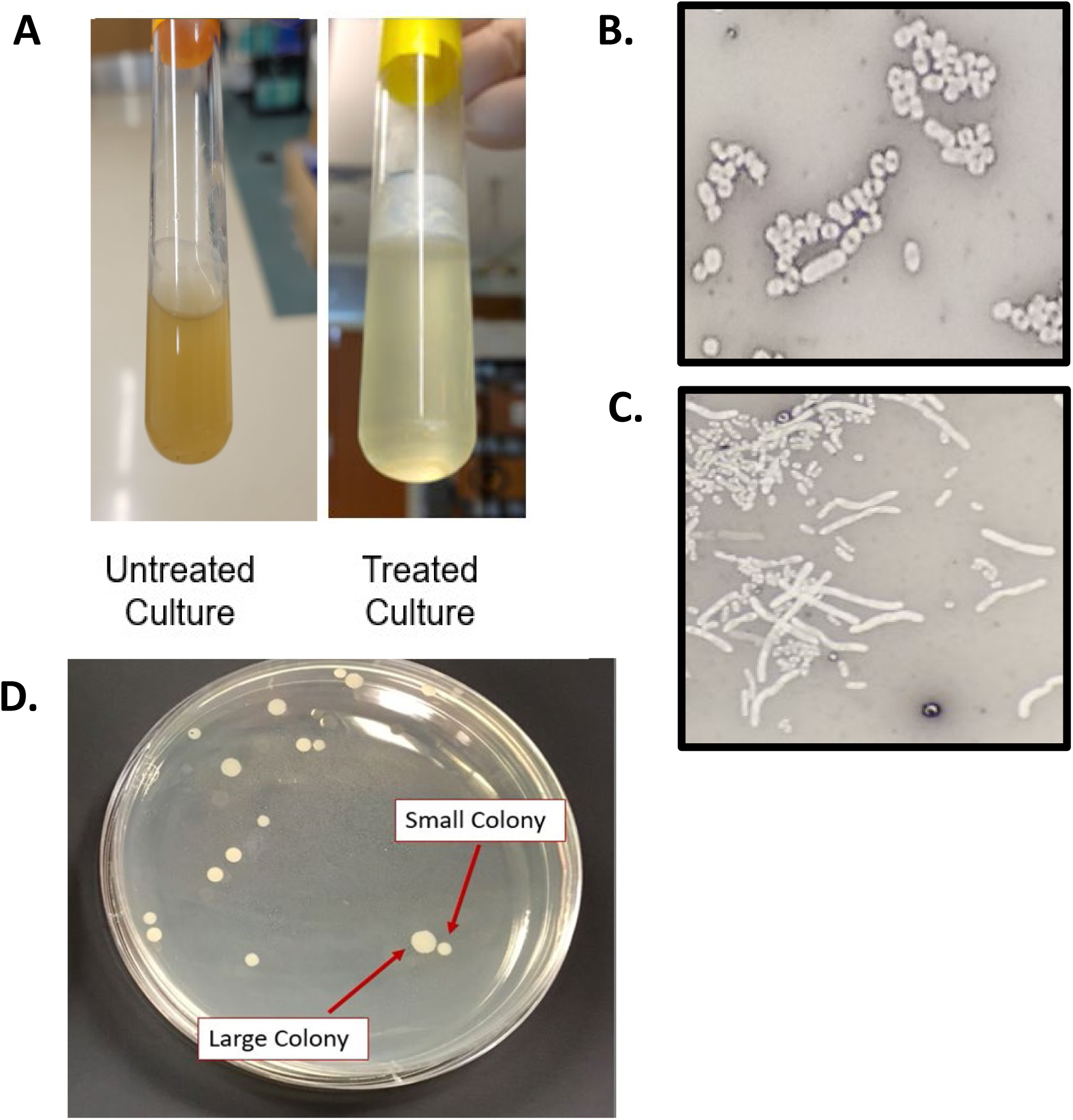
Changes in cell, liquid culture, and colony morphology because of antibiotic exposure. A) Broth culture appearance of untreated and antibiotic exposed culture at end point of the 14-day experiment (after 12 hours of growth) exhibits increased mucoid layer in antibiotic treated culture. Cell morphology of untreated culture (B) and antibiotic exposed culture (C) was determined by negative staining with nigrosin at 1000x magnification. Antibiotic exposed cells (C) exhibit an elongated cell morphology. D) Quadrant steaking on LB of the final adapted culture revealed two colony morphologies.

Differences were also observed in cellular and colony morphology. The antibiotic exposed culture resulted in cells with a greatly elongated, filamentous structure (Figures 2B and 2C). Quadrant streaking of the final antibiotic adapted culture resulted in two consistent colony morphologies, designated small and large colonies (Figure 2D). Bacteria were isolated from either the large or small colonies and resuspended in liquid culture. When replated separately, they only produced small or large colonies that corresponded to morphology of the original colony (data not shown). The persistence of these phenotypes after isolation indicated that the morphological change was heritable.

### Progressive antibiotic exposure induces clinical resistance to multiple classes of antibiotics

Changes in the minimum inhibitory concentration (MIC) of several different classes of antibiotic were determined by the broth microdilution assay (Table 2). The concentration of antibiotic that halts all growth (MIC breakpoint) was determined for tetracycline, amikacin, and five different beta-lactam antibiotics (Cephalothin, Cefoxitin, Cefotaxime, Cefepime, and Imipenem). The beta-lactam drugs were chosen as representative of different generations of cephalosporin antibiotics and a carbapenem, respectively. Clinical resistance was determined using CLSI breakpoints (27).

**Table 2:**
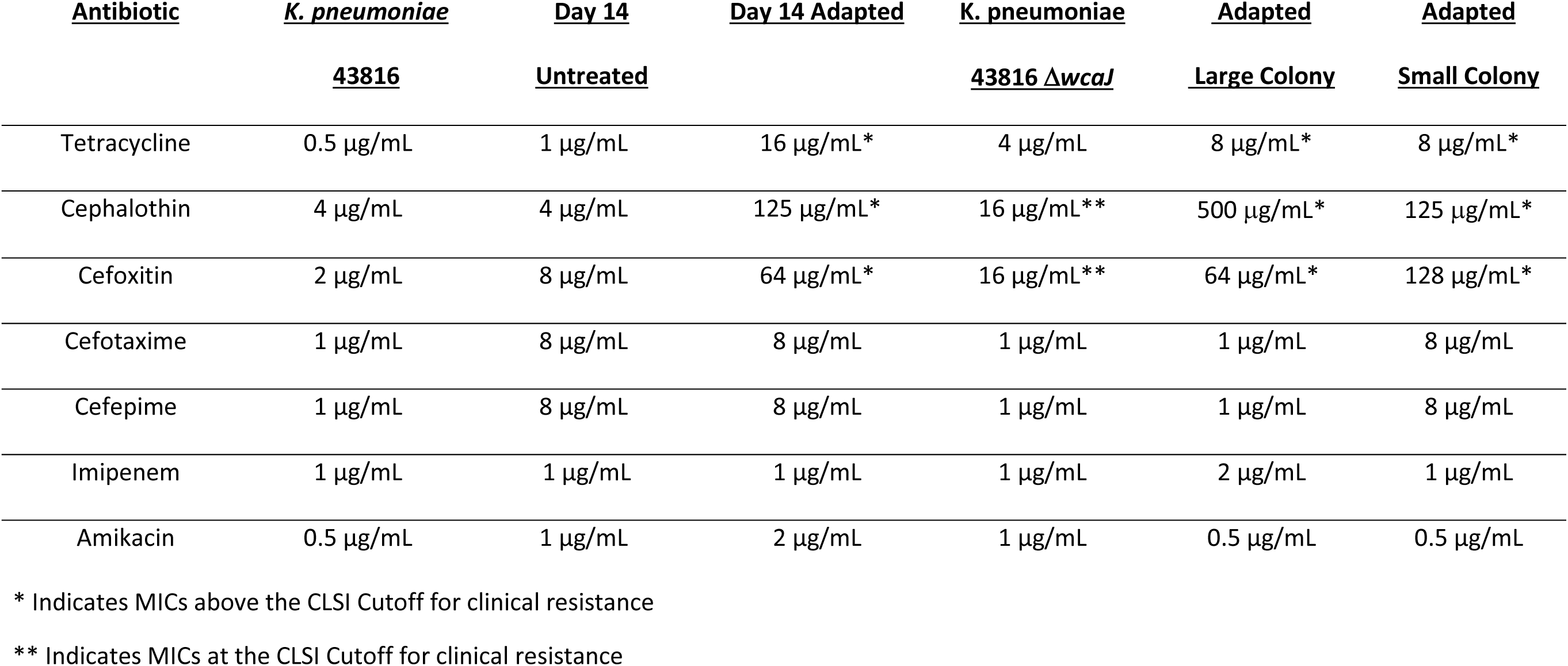
Changes in minimum inhibitory concentration (MIC) breakpoints of *K. pneumoniae* 43816 cultures after progressive low dose exposures. Broth microdilution assays (n=3) determined the minimum inhibitory concentration breakpoint of multiple antibiotics for endpoint cultures, the Δ*wcaJ* strain, and endpoint colony morphology isolates. * indicates values that exceed the CLSI breakpoint for clinical resistance. ** indicates values that meet the CLSI breakpoint for clinical resistance (27).

| <u>Antibiotic</u> | <u>*K. pneumoniae*</u><br><u>43816</u> | <u>Day 14</u><br><u>Untreated</u> | <u>Day 14 Adapted</u> | <u>*K. pneumoniae*</u><br><u>43816 $\Delta wcaJ$</u> | <u>Adapted</u><br><u>Large Colony</u> | <u>Adapted</u><br><u>Small Colony</u> |
| --- | --- | --- | --- | --- | --- | --- |
| Tetracycline | 0.5 µg/mL | 1 µg/mL | 16 µg/mL* | 4 µg/mL | 8 µg/mL* | 8 µg/mL* |
| Cephalothin | 4 µg/mL | 4 µg/mL | 125 µg/mL* | 16 µg/mL** | 500 µg/mL* | 125 µg/mL* |
| Cefoxitin | 2 µg/mL | 8 µg/mL | 64 µg/mL* | 16 µg/mL** | 64 µg/mL* | 128 µg/mL* |
| Cefotaxime | 1 µg/mL | 8 µg/mL | 8 µg/mL | 1 µg/mL | 1 µg/mL | 8 µg/mL |
| Cefepime | 1 µg/mL | 8 µg/mL | 8 µg/mL | 1 µg/mL | 1 µg/mL | 8 µg/mL |
| Imipenem | 1 µg/mL | 1 µg/mL | 1 µg/mL | 1 µg/mL | 2 µg/mL | 1 µg/mL |
| Amikacin | 0.5 µg/mL | 1 µg/mL | 2 µg/mL | 1 µg/mL | 0.5 µg/mL | 0.5 µg/mL |

\* Indicates MICs above the CLSI Cutoff for clinical resistance
\*\* Indicates MICs at the CLSI Cutoff for clinical resistance

As seen in Table 2, the fourteen-day growth of cells without antibiotics did not greatly impact the MIC values against the antibiotic tested. The MIC values in the fourteen-day untreated cells were not as substantially elevated as the MIC values observed in the fourteen-day adapted cells when compared to the day 0 control (Table 2). Overall, the untreated cells did not reach MICs above the CLSI cutoff for clinical resistance for any of the antibiotics tested unlike the fourteen-day adapted cells. The adapted culture exhibited highly elevated MICs to cephalothin, which was used for the progressive exposure experiment. This culture exhibited clinical resistance to cephalothin and cefoxitin as determined by CLSI breakpoints. Cephalothin and cefoxitin are first- and second-generation cephalosporin antibiotics, respectively. MICs to later generation cephalosporins and the carbapenem antibiotic were slightly elevated in both the fourteen-day untreated and adapted cells, but the MICs for each were still below the CLSI cutoff. To fully evaluate the impact of progressive sub-MIC cephalothin exposure, antibiotics with different cellular targets than the beta-lactams were also tested. Progressive exposure had little impact on the breakpoint value for the aminoglycoside antibiotic amikacin, but clinical resistance to tetracycline was achieved. These data indicate that progressive exposure to cephalothin at only half the MIC cutoff concentration (27), led to resistance to a second generation of cephalosporin in addition to an antibiotic in another class.

Additionally, MIC breakpoints were determined for the two isolated colony morphologies from the final antibiotic adapted culture. Both the small and large morphologies exhibited clinical resistance to cephalothin and cefoxitin, with the large colonies exhibiting highly elevated MICs to cephalothin; which was well above the MIC observed for both the small and adapted final culture. Both isolates also exhibited elevated MICs to tetracycline; but were more sensitive than the mixed day 14 adapted culture. Amikacin MICs were not significantly changed in either phenotype. Taken together, these data demonstrate that progressive sub-MIC concentrations of a single antibiotic provide sufficient evolutionary pressure to drive the evolution of clinical levels of resistance to multiple generations of beta-lactams as well as classes of antibiotics with different cellular targets.

### Whole genome sequencing reveals distinctive genetic changes because of progressive antibiotic exposure

To determine the genetic changes responsible for this antibiotic resistant mucoid phenotype, DNA was extracted from bacterial stocks of each day of the progressive antibiotic exposure and sequenced by the Microbial Genome Sequencing Center using an Illumina sequencing method and a proprietary statistical algorithm for variant calling (www.seqcenter.com) (16). This methodology resulted in a surprisingly low number of SNPs identified, in part because it does not analyze changes to non-coding regions of the genome, such as promoters.

To confirm these results and to extend our analysis to include SNPs within promoter regions, DNA from the fourteen-day adapted and untreated *K. pneumoniae* 43816 cultures were also analyzed using SMRT Cell Sequencing by the National Center for Genome Resources (www.ncgr.org). Together these analyses identified 107 mutations in the control culture and 29 mutations in the genome of the antibiotic treated culture. Of the 29 mutations identified in the adapted strain, 15 were nonsynonymous changes that occurred in protein-coding regions of the sequence. To further refine the list of potential targets, any mutations shared with the untreated population of bacteria were removed. Comparing the genomes highlighted seven protein coding genes which were altered in the final antibiotic adapted cells (Table 3). Changes to *rpoB*, *tetR/acrR*, *wcaJ*, and *gndA* were identified using both sequencing methodologies (Table 4).

**Table 3:** Non-synonymous single nucleotide polymorphisms and genomic changes identified in the final adapted culture. SMRT Cell genomic sequencing was used for whole genome sequencing. The genome sequence of *K. pneumoniae* 43816 was used as the reference genome.

| Position | Change in Sequence | Depth of Coverage | Gene/Promoter | Gene Family |
| --- | --- | --- | --- | --- |
| 10279 | G→C | 43 | Gene | SGNH/GDSL hydrolase |
| 10285 | G→C | 43 | Gene | SGNH/GDSL hydrolase |
| 10412 | T Deletion | 56 | Promoter | SGNH/GDSL hydrolase |
| 412527 | G Deletion | 100 | Gene | N-acetyltransferase |
| 1125537 | Insertion T | 100 | Gene | Globin |
| 2008934 | Insertion TTTCGCTA | 100 | Gene | TetR/AcrR Transcription regulator |
| 2063994 | C Deletion | 100 | Gene | ABC Transporter |
| 3225592 | C Deletion | 100 | Gene | RmuC DNA Recombination |
| 5188116 | T → G | 100 | Gene | UDP Phosphate Glucose Phosphotransferase |
| 2657272 | G Deletion | 100 | Promoter | Peptide Chain Release Factor 3 |
| 1078904 | C Deletion | 100 | Promoter | LysR Transcriptional Regulator |
| 1294495 | G Deletion | 100 | Promoter | Type II Asparaginase |
| 1823692 | A Insertion | 100 | Promoter | Pyrimidine Photo-lyase |
| 5050494 | A Insertion | 100 | Promoter | Cyclic Di-GMP Phosphodiesterase |

**Table 4:** SNPs and genomic changes identified in large and small colony subpopulations of 14 day adapted culture. Illumina genomic sequencing was used for sequencing of colony subpopulations. The genome sequence of *K. pneumoniae* 43816 was used as the reference genome.

| Colony Variant | Position | Original | New | Coverage<br>(Small/Large) | Gene |
| --- | --- | --- | --- | --- | --- |
| Small and Large | 2,008,948 | . | Insert TATTTCGC | 100/52 | TetR/AcrR family transcriptional regulator |
| Small and Large | 3,191,984 | A | T | 100/100 | *rpoB*/DA-directed RNA polymerase subunit beta |
| Large | 1,548,640 | Unspecified | Unspecified | 15 | ComEC family protein |
| Small | 4,676,685 | T | G | 100 | YfiR family protein |
| Small | 5,187,773 | Deletion 2,103 bp. | . | 100 | *wcaJ* and *gndA* |
| Final adapted mixture | 5,187,772 | T | G | 64 | *wcaJ* Undecaprenyl phosphate-glucose phosphotransferase |
| Final adapted mixture | 5,189,876 | unspecified | unspecified | 64 | *gndA* NADP-dependent phosphogluconate dehydrogenase |

Notably, none of the identified mutations by either methodology were identified in penicillin binding proteins or other cell wall modification genes highly associated with traditional beta-lactam resistance. Genes with identified mutations were searched against the CARD data base and using the *amrFinder* bioinformatics tool (card.mcmaster.ca) (28)(29). Neither of these analyses identified these genes as being previously identified as resistance mechanisms to beta-lactams. The *tetR/acrR* gene resulted in a match for the wildtype version of the gene as being associated with tetracycline resistance, while *rpoB* matched for rifamycin resistance (30). This analysis indicates that the majority of these modifications and gene targets are novel or not previously characterized in terms of their impact on antibiotic resistance.

The only mutations that were not 100% fixed in the population at the conclusion of the fourteen-day progressive exposure were nucleotide substitutions in the coding region of the SGNH/GDSL hydrolase family protein and a deletion in the promoter of SGNH/GDSL (Table 3). This may be linked to the emergence of the small and large colony variants. Other genetic changes that were fixed by the fourteen day endpoint include deletions in the coding regions of the following: N-acetyltransferase, ABC transporter, ATP binding protein, and DNA recombination protein (RmuC). Additionally, a substitution in the coding region of undecaprenyl-phosphate glucose phosphotransferase was discovered. An insertion was found in a globin coding sequence and a large insert in the *tetR/acrR* transcriptional regulator resulted in an early stop codon. The functions of these genes encompassed cellular processes of signal transduction, energy and metabolite use, capsule formation, and nucleic acid proofreading.

In addition to changes in the coding regions of those genes, six unique mutations were identified in promoter regions of five other genes that were fixed in the adapted population (Table 3). Promoter regions were defined as occurring within 150 bp of a protein coding start site. Synonymous mutations between the untreated and adapted cultures were removed as described above. With exception to SGNH/GDSL, all other genetic changes in promoter regions were fully fixed in 100% of the DNA sampled. Not all genes associated with these promoters have been fully characterized in *Klebsiella pneumoniae.* In those cases, the data about the class of each gene and close homologues is presented. Functions of the coding regions encompassed cell metabolism, signal transduction, and mRNA proofreading.

### Genome sequencing allows for correlation of genetic changes with increased MICs

The Illumina sequencing of the cultures from each day of the experiment allowed us to map the timepoint of 100% fixation of mutation with significant increases in MIC, and changes in growth rates. For this we focused on those mutated genes (*rpoB*, *tetR/acrR*, *wcaJ*, and *gndA)* identified using both sequencing methodologies (Table 4).

As seen in Figure 3, the emergence of clinical tetracycline resistance can be matched with fully fixed alterations in *rpoB*, and large increases in the cephalothin MICs can be correlated with fixed changes to *tetR/acrR*, and *wcaJ.* The large increase in cephalothin MIC associated with *wcaJ* can also be mapped to the significant slowing of growth of all treated cultures when exposed to 5 μg/ml of drug (Table 1).

**Figure 3:**
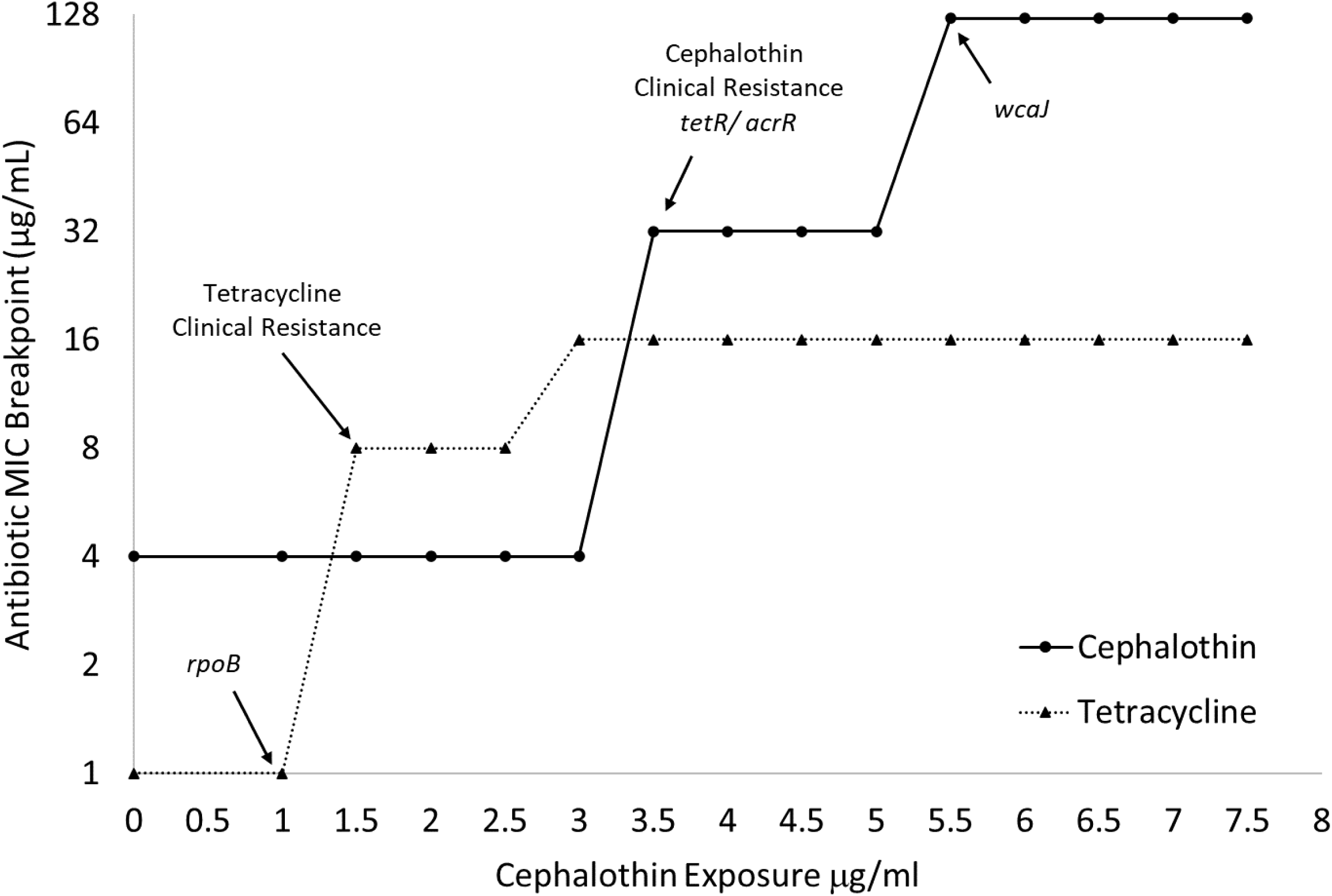
MIC breakpoints map to onset of clinical resistance and fixation of genomic changes. Broth microdilution assays (n=3) determined MIC breakpoints to tetracycline and cephalothin of cultures over the entire 14-day progressive exposure. Arrows indicate jumps above clinical resistance breakpoint and fixation of specific genetic changes as identified by whole genome sequencing.

Two separate sets of sequence changes in *wcaJ*, and *gndA* were identified using the Illumina methodology (Table 4). The final adapted culture, which is composed of a mix of large and small colony morphologies, identified sequence changes with only 64% genome coverage. When the two morphologies were sequenced separately, a large deletion that impacted both genes was found in only the small colony variant.

Both the large and small colony variants had mutations in the *rpoB* and *tetR/acrR* transcriptional regulator sequences. The mutation in *rpoB* was 100% fixed in both variants. However, the depth of coverage of the *tetR/acrR* transcriptional regulator in the large colony variant was only 52%. The large colony variant also had a distinguishing marginal mutation call in a c*omEC* family protein that was not found in the small colony variant. The small colony variant exhibited an additional SNP in a *yfiR* family protein that was not identified in the large colony variant. All the mutations in the small colony forming variant were 100% fixed indicating a more homogenous genomic identity than the large colony forming subpopulation.

### Genetic changes in progressive antibiotic adapted cultures are associated with alterations to capsule and LPS

The final adapted culture from this progressive antibiotic exposure experiment exhibited a highly mucoid phenotype indicating that capsule production or composition may be altered. Genome sequencing then identified alterations to *wcaJ*, which is part of the capsule *cps* operon and initiates production of colanic acid (31). Therefore, we hypothesized that alterations to capsule production and composition may be directly related to the emergence of high concentration cephalothin resistance.

To investigate changes in the capsule, extracellular polysaccharide was extracted and quantified by the uronic acid assay. As can be seen in Figure 4A, capsule polysaccharide increases until the fixation of the *tetR/acrR* mutation and emergence of clinical cephalothin resistance. All samples after this point exhibited capsule polysaccharides at levels similar to or slightly below those produced by the untreated control culture.

**Figure 4:**
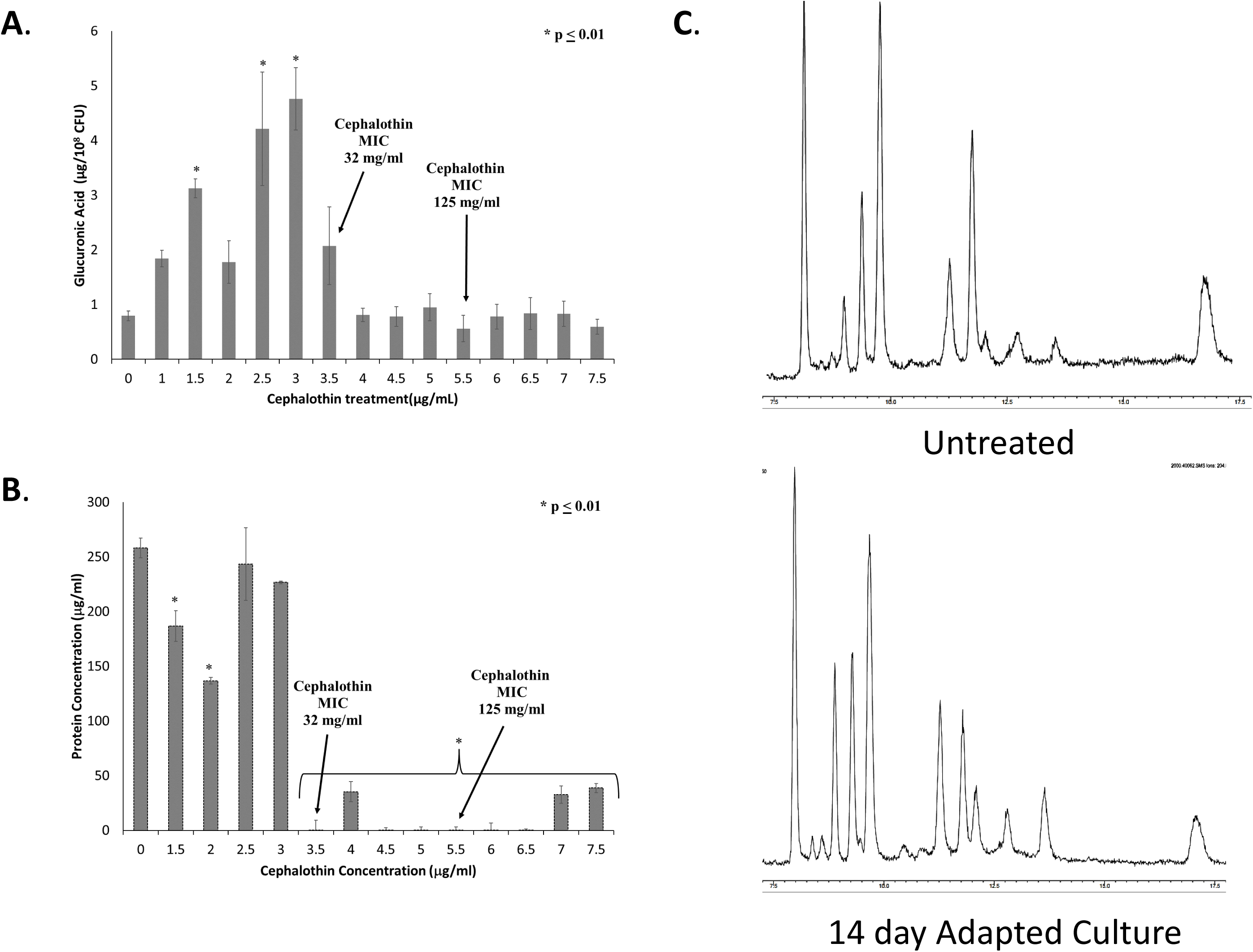
Onset of clinical resistance coincides with changes in quantity and composition of capsule. Capsule polysaccharides were extracted from cultures from each treatment of the experiment. The extracted capsule was then analyzed for A) Total capsule production as quantified by glucuronic acid; and B) Protein content of extracted capsule quantified by BCA protein assay. N=6 for both assays, *= ≤ 0.01 by one way ANOVA compared to untreated control. C) GC-MS analysis of the carbohydrate composition of capsules extracted from untreated and adapted cultures.

Given that the uronic acid assay only detects one sugar moiety within the capsule polysaccharide, other possible components of the capsule were investigated. Extracted capsule from untreated and adapted cultures were found to contain similar levels of sialic acids (2.5 ± 1.2 nmoles/10^8^ CFU in untreated culture vs 2.9 ± 1.8 nmoles/10^8^ CFU in final adapted culture), which is associated with increased virulence (32). Adapted capsule extracts also exhibited reduced total protein content as determined by BCA assay (Figure 4B). Both the reduced glucuronic acid and protein emerged concurrently with the alterations in the *tetR/acrR* gene. Finally, capsule extract was analyzed by GC/MS to identify new peaks indicative of changes to the sugar composition (Figure 4C). No new peaks were identified indicating that novel carbohydrate changes are not directly associated with the antibiotic resistant phenotype.

To determine the relationship between alterations in capsule and antibiotic resistance, we tested a Δ*wcaJ* strain of *Klebsiella pneumoniae* 43816 for altered MICs. This deletion has been previously characterized as significantly reducing capsule uronic acid content and mucoviscosity (12). As seen in Table 2, deletion of this gene does result in increases in MIC breakpoints to both cephalothin and cefoxitin. These increases are to the level of the breakpoint value. This data indicates that cps modification can have a significant impact on antibiotic susceptibility in *Klebsiell*a. However, the high-level clinical resistance seen in our fully adapted strain is likely due to the combination of multiple gene alterations, including those within the cps operon.

Colanic acid, which is synthesized in part using the *wcaJ* gene product, is also associated with lipopolysaccharide. Therefore, we determined the LPS content of both bacterial cells and extracted capsule using the Purpald assay. LPS content was significantly elevated in both the cell and extracted capsule of the adapted strain as compared to the control (Figure 5). This elevated LPS content may indicate a high level of outer membrane turnover in the adapted culture associated with alteration in *wcaJ*.

**Figure 5:**
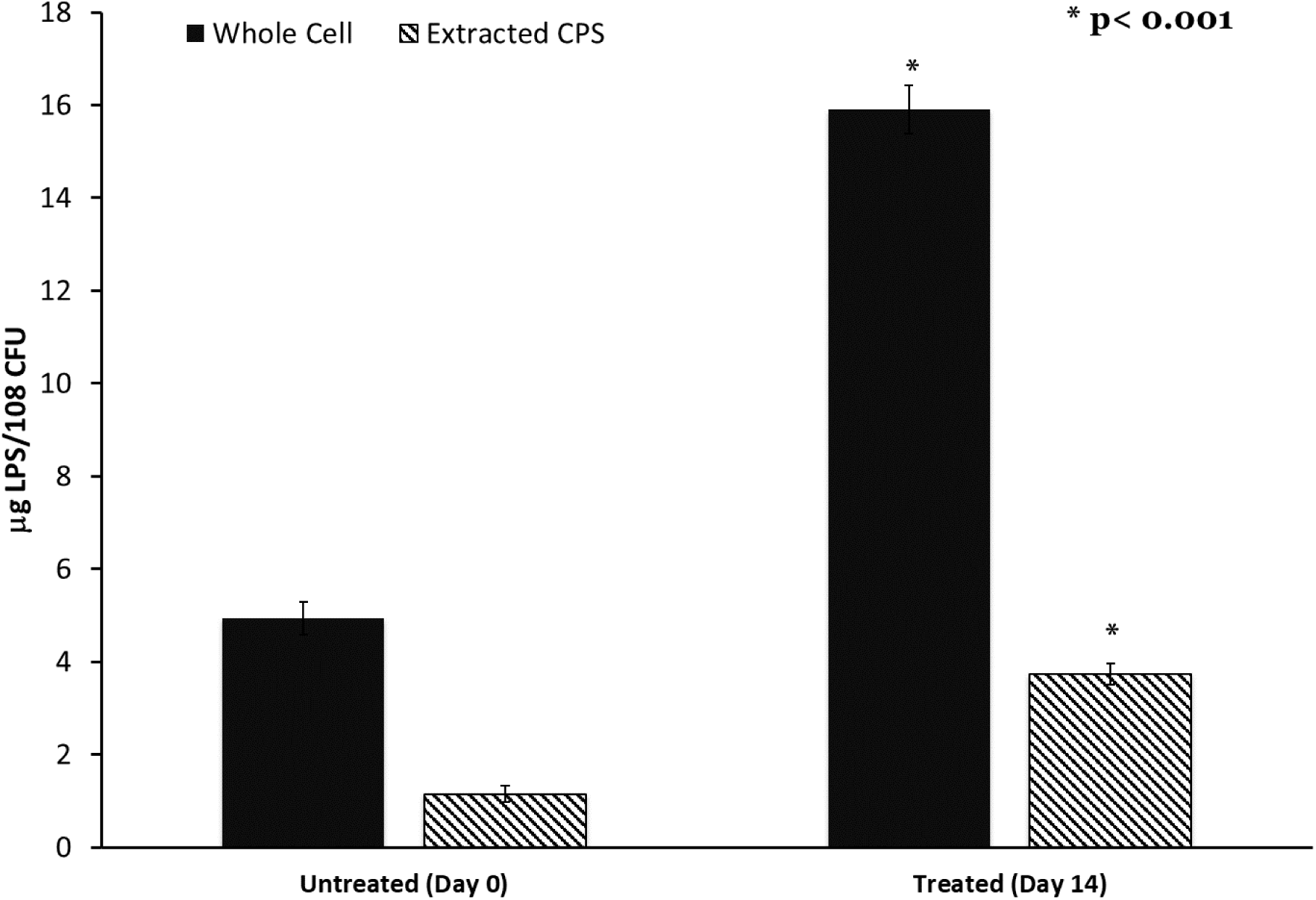
Lipopolysaccharide content of capsule and cells is increased after progressive sub-MIC exposure. Capsule polysaccharides were extracted from untreated and final adapted cultures. Capsule and whole cells LPS content was determined by the Purpald Assay. N=6, *= ≤ 0.001 by one way ANOVA compared to untreated control.

## Discussion

The goal of this study was to characterize the impact of low concentration antibiotic exposure on the evolution of bacterial resistance. A progressive exposure model was used, in which *Klebsiella pneumoniae* was exposed to slowly increasing sub-MIC concentrations of the antibiotic cephalothin. The final resulting culture had been exposed to a maximum antibiotic concentration of 7.5 μg/mL cephalothin. This final culture exhibited full clinical resistance to first and second generation cephalosporins, with MIC values far exceeding the CLSI breakpoint. Additionally, the final culture had increased MICs to tetracycline, a highly mucoid appearance, and an elongated cellular morphology. Genome sequencing revealed a series of genetic alterations that could be mapped directly to the emergence of the resistant phenotype. Changes in phenotype and genetic alterations both indicate that alterations in the *tetR* regulator, LPS shedding, and capsule colanic acid synthesis may be directly associated with the resistance phenotype.

The three resistance-correlated mutations occurred in genes *rpoB*, *tetR/acrR*, and *wcaJ*. The increases in resistance occurred in a stepwise manner, similar to the increases in fitness observed in *E. coli* long-term evolution (LTEE) experiments (33–35). LTEE studies also provided estimates for the rapidity of mutation fixation in a constant environment. The *E. coli* population in the LTEE experiments accumulated 20 mutations in the first 10,000 generations of growth with a few rapid mutations that reached fixation in the population within 100 generations (34, 35). The three resistance-correlated mutations in this present study all fixed within 36-378.9 generations using a generation time estimate from a related *K. pneumoniae* strain of 38-40 minutes (36). Despite the use of incremental exposures to cephalothin in this study, we find that these mutations occurred rapidly in *Klebsiella pneumoniae* similar to what others have reported for *E. coli*.

The speed with which these three mutations were acquired also indicates that the sub-MIC concentration of cephalothin in the growth environment provides significant selective pressure. After only 24 hours of exposure, the resistance of the adapted population to tetracycline had the first jump from 1 µg/mL to 8 µg/mL, which correlated with a fixed mutation in *rpoB*. The rapidity of fixation and correlated increase in survivability suggest that this mutation conferred a very high fitness advantage. It is possible that a necessary step in acquiring resistance for this population was mutation in a gene meant to monitor nucleic acid integrity, allowing for further alterations to the genome.

The selection window hypothesis holds that sub-MIC level antibiotic exposure should generally increase bacterial mutation rates (7, 8) because higher mutation rates will improve the chances of generating adaptive genetic change that provides a fitness advantage in conditions of antibiotic stress. Methodologies employed by studies testing this theory have generally identified “mutants” by plating samples of a bacteria exposed to an antibiotic on media inoculated with some other antibiotic substance (4, 9). Mutation rates analyzed using this methodology tend to indicate that lower antibiotic concentrations result in high mutation rates that decline as the concentration of antibiotic nears the MIC (9, 37). However, when whole-genome analysis is incorporated along with mutation accumulation analysis, substitutions and insertions/deletions increased in frequency as exposure to antibiotic increased in *E. coli* (38). Additionally, the rate of mutation in known resistance genes tracked in *P. aeruginosa* isolates generated conflicting results (11).

The present study utilized whole genome sequencing to identify mutations across the genome for the antibiotic-treated and unexposed sample after 14 days of total treatment. This data showed a higher number of mutations in the untreated sample compared to the adapted cohort. As noted by Long et. al., a true mutation accumulation analysis that determines the frequency of genetic polymorphism per generation would require repeatedly passing a bacteria through bottlenecks to mitigate any selective influences (38). The fact that the adapted population in this study was maintained as a whole group when sub-cultured may affect the relative number of mutations identified by whole genome sequencing. Another factor at work is that the treatment concentration of cephalothin did surpass the MIC of 4.0 μg/ml of the original *K. pneumoniae* 43816 for over half of the adaptation period (Tables 1 and 2). Both the selection window hypothesis and adaptive studies of bacterial mutation would suggest that such a high concentration would decrease mutation rates compared to an unexposed cohort and explain the low number of identified polymorphisms (4, 7–9, 37).

The *rpoB* mutation identified by Illumina sequencing early in the exposure regimen is very close to an alteration detected in the *rmuC* DNA recombination protein in the final sample sequenced using the SMRT cell method. The mutations identified do not all match between both sequencing methods, nor are the calls for similar genes found in the exact same position or using the same base changes (Tables 3 and 4). This could reflect differences in methodology, genome construction in the two methods, and/or inherent error. These results highlight the limitations of using just one sequencing methodology for this type of analysis. Recent work has suggested that both long-read methods, like SMRT cell sequencing, and short-read methods, like Illumina sequencing, be combined for a hybrid approach for the most accurate formation of novel sequences especially those of the *Enterobacteriaceae* family (39). The data from both rounds of sequencing found that the SMRT cell method identified more polymorphisms in the adapted population genome and was able to identify mutations in noncoding regions unlike the Illumina method. Additionally, the evolution of a resistant phenotype can require cooperative mutations adding difficulty to the present study’s ability to directly trace the effects of one mutation to a quantifiable change in antibiotic survival (37).

The protein products of both *rpoB* and *rmuC* interact closely with genomic DNA. RmuC is a regulator which can prevent sequence inversion during replication (40). RmuC has also been identified as a possible multidrug resistance (MDR) gene in other Gram-negative bacterial species (41, 42). However, RpoB is part of the RNA polymerase protein complex and when mutated can inhibit the action of rifamycin in bacteria (30). Mutations in *rpoB* have been linked to rifamycin resistance in *E. coli* (43, 44), and has been identified as a resistance gene in various other classes of bacteria (45). Additionally, an early mutation in a gene tied to nucleic acid integrity provides a possible mechanism for further evolution of resistance by increasing the occurrence of replication error mutations in progeny.

TetR regulators are global multi-target transcriptional regulators that affect multiple processes within the cell beyond just efflux pumps (43, 44). Members of this family of regulatory proteins have been shown to impact a variety of virulence associated targets including motility, biofilm formation, and osmotic tolerance (46). One member of this family has been directly associated with *ftsZ*, which is known to regulate cell division and bacillus cellular morphology (47). Therefore, it is reasonable for a mutation in a *tetR* type regulator to impact cellular functions and resistance to classes of antibiotics other than tetracycline.

Other studies on the *in vivo* evolution of *Klebsiella* within a patient undergoing antibiotic therapy have further identified TetR family upstream regulators of porins as being associated with clinical resistance. Yoshino et. al, tracked the evolution of a clinical isolate of *Klebsiella* through treatment with multiple classes of antibiotics. They identified porin loss as a result of mutations in the *ramR*, a TetR family transcriptional regulator of *micF*, the an antisense RNA regulator of the ompK35 porin (48). The ultimate result of this *ramR* mutation was loss of the OmpK35 porin, limiting entry of beta-lactam antibiotics into the periplasmic space. These data indicate that antibiotic resistance can be achieved through mutations in a number of different genes in a given pathway. In a sub-MIC exposure to drug, it may even be advantageous for mutations to occur in regulator proteins rather than negatively impacting the primary function of a porin or another endpoint enzyme.

None of the identified mutations were found in penicillin binding proteins or other cell wall modification pathways that are highly associated with traditional beta-lactam resistance. The data presented here adds to a number of other recent studies that are identifying the role of a broad array of genes and cellular functions as being involved in the bacterial response and survival during antibiotic exposure. In addition to regulators of porins and cell wall regulators, Lopatkin *et. al* have identified core metabolic mutations that mitigate antibiotic susceptibility in clinical isolates (49). Together these findings all support the need for broader investigations of gene alterations that contribute to resistance than those traditionally classified as such.

It is interesting that such a large insert and formation of a stop codon would be found in *tetR/acrR,* a gene known for resistance to tetracyclines, when exposed to a beta-lactam. However, the development of cross resistance among bacterial populations exposed to one class of antibiotic is not uncommon. Exposure to environmental chemicals such as surface antiseptic chlorhexidine, or veterinary antibiotics tilmicosin and florfenicol have been shown to create cross-resistance in human pathogens to different classes of antibiotics (3, 5).

The genetic alterations associated with the largest increase in beta-lactam resistance were mapped to two adjacent genomic locations. The SMRT method identified an uncharacterized undecaprenyl phosphate-glucose phosphotransferase, which is very close to the mutation found in *wcaJ* by Illumina sequencing (Tables 3 and 4). The second round of sequencing specified that there was a large deletion encompassing the end of the *wcaJ* and beginning of the *gndA* genes which was exclusive to the small colony forming subpopulation.

In *Klebsiella pneumoniae*, *wcaJ* is part of the capsular *cps* operon and initiates production of colanic acid (31, 50). Absence of *wcaJ* has been linked to increased resistance to phage treatment, decreased virulence in murine models, and increased phagocytosis by macrophages (31). Studies ablating *wcaJ* in *Klebsiella pneumoniae* result in a nonmucoid phenotype while increasing biofilm production and increasing resistance to polymyxin (12)(50).

Our observation of altered cellular morphology may be related to envelope stress and remodeling involving enzymes such as WcaJ. This elongated morphology has been demonstrated in experiments using high doses of beta-lactams to be a signature event in the process of drug induced cellular lysis (51). However, other studies of beta-lactam exposure have documented the permanent formation of filamentous bacterial cells (52). Additionally, Kessler *et. al* demonstrated altered cell shape in *E. coli wcaJ* knockouts that exhibited accumulation of periplasmic colanic acid precursors and activation of the Rcs osmotic stress response pathway (53). New studies indicate that changes in cellular shape and the surface area to volume ratio can be part of an overall bacterial stress response to antibiotics (54). Our results, which only used sub-MIC concentrations of beta-lactams, indicate that these stress responses alone can result in permanent changes to cellular morphology that may play a larger role in bacterial survival.

Colanic acid polymers, synthesized using the *wcaJ* gene product, can be covalently linked to lipopolysaccharides (55). We observed increased LPS content in our extracted capsule polysaccharides, indicating that the *wcaJ* mutation may trigger increased membrane turnover. The ability to synthesize LPS O-antigen sugars has been documented to directly impact colanic acid synthesis (56). It is therefore reasonable to hypothesize that alterations in *wcaJ* may likewise impact LPS synthesis, modification, and turnover. Studies of more general osmotic stress indicate the possibility of a shift from O-antigen attachment to colanic acid attachment (55). These changes may directly impact overall membrane permeability and the ability of beta-lactams to access the periplasmic space.

At low antibiotic concentrations, these modifications in membrane composition and permeability may be sufficient for bacterial survival. This idea is supported by our analysis of MIC breakpoints to cephalosporins from multiple different generations. The multiple generations of cephalosporin antibiotics are classified in large part based on molecular modifications that extend the spectrum of activity against gram-negative aerobic bacilli such as *Klebsiella* (57). We observed increased MICs to first and second, but not later generation cephalosporins in both our adapted strain and the Δ*wcaJ* strain. These later generation drugs, as a well as carbapenems have been modified to enhance permeability through the Gram-negative outer membrane. These data indicate that the mutations generated by our adapted stain may have directly impacted outer membrane permeability by a mechanism other than porin loss.

Together, these data indicate that low level beta-lactam exposure initiates a cascade of modifications to the outer envelope and capsule polysaccharides that warrants further investigation. Studies examining single knockouts of *tetR* did not find associated changes in beta-lactam MICs (46). While we observed increases in the MICs in the Δ*wcaJ* strain of *Klebsiella pneumoniae* 43816, Kessler et al. did not when using a knockout of the same gene in *E. coli* (53). This may be related to the more complex nature and function of the capsule in *Klebsiella*. Mutations in *rpoB* have been associated with beta-lactam resistance, but not resistance to multiple classes of antibiotics (58). It is therefore highly likely that a combination of multiple mutations is required to achieve the multi-drug resistant phenotype observed in this experiment.

Our final adapted culture represented a heterogeneous mixture of two colony morphologies, which was reflected in the genome sequence analysis. The deletion spanning between *wcaJ* and *gndA* only occurred in the small colony forming population. As noted above, ablation of *wcaJ* in *K. pneumoniae* is linked to a decrease in species mucoidy and increased sensitivity to polymyxin. Similarly, *gndA* is a gene within the *cps* locus responsible for the K2 serotype and capsule formation of *K. pneumoniae* (59, 60). It therefore seems likely that the mucoid phenotype seen in the fourteen-day adapted population whole-group mixture is due to the large colony subpopulation only. This is supported by the fact that the small colony subpopulation has a reduced resistance to cephalothin compared to the large colonies which have an intact *wcaJ* sequence.

This study also demonstrates the complexity of changes occurring within what can be characterized as a single culture study. Multiple mutations occur within different cells in this population over time, resulting in a heterogeneous, non-clonal population. As compared to studies such as the LTEE (33–35), we have added the directional pressure of sub-MIC antibiotic exposure, and prevented the acquisition of new resistance genes from other nearby bacterial species, such as might be found with a patient. Tracking the *in vivo* evolution of clinical isolates has been used to demonstrate the evolution of resistance by a single *Klebsiella* isolate to colistin (61), carbapenems (62) and beta-lactams (48, 63). However, these studies often focus primarily on the acquisition of genes by horizontal gene transfer, and only recently have identified mutations in other pathways as playing a significant role (54)(49) in bacterial survival within the antibiotic treated host. Larger scale studies, using multiple cultures provided the same antibiotic exposure in parallel are ongoing to investigate how many different paths and mutations may be used to ensure bacterial survival.

In summary, this progressive, sub-MIC antibiotic exposure experiment resulted in a mixture of bacteria exhibiting multiple genomic changes. The multiple isolates from this experiment resulted in bacteria that were resistant, rather than tolerant of the antibiotic, as they exhibited normal metabolic activity and growth in the presence of the antibiotic (64). While both the large and small variant isolates exhibited elevated MICs to cephalothin, the mixture of these isolates demonstrated a synergistic protective effect. This mixture is reflective of what would occur in the environment, where individual cells may independently evolve, persist, or assist in the survival of nearby cells. The bacteria in this study achieved clinical resistance without the traditional acquisition of a beta-lactamase gene. These compensatory mutations, and combinations of them, warrant further investigation as they may accelerate and enhance resistance associated with traditional horizontal gene transfer.

## Funding and Acknowledgements

This research was supported by funding from UNF to T.N.E., including a Dean’s Council Fellowship Award, a Research Enhancement Plan Award, and the Transformational Learning Opportunity Program. We thank Frank Smith for assistance with bioinformatics software, and all members of the Ellis lab for thoughtful input and comments.

## Notes

### Competing Interest Statement

The authors have declared no competing interest.

### Summary of Updates

This revision addressed reviewer concerns. Major additions include analysis of wcaJ knockout strain and expanded analysis of multiple generations of cephalosporin antibiotics.

